# Arabidopsis histone demethylase RSI1/FLD self-regulates its pathogen-induced expression

**DOI:** 10.64898/2026.11.19.767995

**Authors:** Uday Singh, Vishal Patil, Ranjan Kumar, Vijayata Singh, Ashis Kumar Nandi

## Abstract

REDUCED SYSTEMIC IMMUNITY 1(RSI1)/FLOWERING LOCUS D (FLD) is essential for the activation of systemic acquired resistance (SAR). RSI1 functions as a repressor of gene expression by demethylating H3K4me2 and H3K4me3. *RSI1* expression is induced both in local and systemic tissues upon pathogen inoculation. However, the mechanism of its induction or its expression regulation upon pathogenesis is not known. Here we show that *RSI1* expression in SAR-induced *rsi1* mutant plants is higher than WT plants, suggesting self-regulation. Pathogen-induced expression of RSI1 requires the function of GSTT2, a SANT domain-containing protein essential for SAR activation. Co-expression of RSI1 suppresses pRSI1:GUS activity in a transient assay system. Chromatin-immunoprecipitation (ChIP) experiments reveal that RSI1 physically associates near the transcription start site (TSS) of the *RSI1* locus and facilitates demethylation of H3K4me2 and H3K4me3 at *RSI1* locus. Our data altogether demonstrate that RSI1 negatively regulates its own expression as a possible mechanism of keep the SAR expression at an optimal level.

## 1. Introduction

Plants have a strong defense system to counter a large variety of pathogens. Upon pathogen infection, plants produce a large variety of antimicrobial compounds which help to keep the growth of pathogens under control (Jones & Dangl, 2006; Spoel & Dong, 2012). Production of these antimicrobial compounds is primarily regulated at the transcriptional level of their biosynthetic genes. An experienced plant exhibits stronger defense than a naïve plant through the activation of systemic acquired resistance (SAR) (Adam et al., 2018; Jung et al., 2009). Experiments suggest that in addition to antimicrobial compounds, pathogen-infested tissues synthesize a group of molecules that travel through the phloem to the rest of the plant and function as SAR-mobile signals. Genetic and biochemical experiments identified several SAR mobile signals which include methyl-salicylate (MeSA), dehydroabietinal (DA), glycerol-3-phospahte (G3P), azelaic acid (AzA), pipecolic acid (PiP), extra-cellular ATP (eATP), DEFECTIVE IN INDUCED RESISTANCE 1 (DIR1) protein and others (Chanda et al., 2011; Dempsey & Klessig, 2012; Li et al., 2023; Luna et al., 2012; Návarová et al., 2012; Navarova et al., 2012). SAR mobile signals influence epigenetic modifications in systemic tissues that facilitate infection memory development and heightened resistance.

Though the nature of SAR mobile signals, their biosynthetic pathways, and their impact on SAR are relatively well established, the mechanism by which these signals generate infection memory is little known. Genetic screening of an EMS-mutagenized Arabidopsis population identified *reduced systemic immunity 1* (*rsi1*), which is specifically defective in the perception of SAR mobile signals in systemic tissues (Singh et al., 2013). The *rsi1* mutation is allelic to the *FLOWERING LOCUS D* (*FLD*) gene (Singh et al., 2013). The *rsi1* bears a point mutation in which a conserved glycine is changed to arginine (G^512^ to R^512^). This change makes FLD/RSI1 non-functional, as evidenced by the mutant phenotypes. Like *fld* mutants, *rsi1* shows late flowering and aerial rosettes. Similarly, the T-DNA insertion knockout mutants like *fld3* and *fld4* are defective in SAR, like that of *rsi1* (Singh et al., 2013).

RSI1 expression increases upon pathogen inoculation in both local and systemic tissues. RSI1 codes for H3K4me2/me3 demethylase and function as suppressor of gene expression (Kumar et al., 2025; Singh et al., 2023; Singh et al., 2014). Here we show that RSI1 protein physically associates at its own promoter and negatively regulates its expression.

## 2. Materials and Methods

### 2.1 Plant growth

Plants were grown at 22°C temperature, and 12h/12h light duration and 80 µE m^-2^s^-1^ intenstity. Humidity was 70%. For plant growth, soil containing a soilrite and vermiculite (6:1) mixture was used. Plants were regularly watered at intervals (Giri et al., 2017; Singh et al., 2013).

### 2.2 GUS Assay

*RSI1* promoter along with GUS was cloned into the pCambia1302 vector (pRSI1:GUS), and RSI1 CDS into pCSXN (p35S:RSI1-HA). pCambia1302 empty vector and pCSXN-HA empty vector (p35S:HA) was used as control. Further, these constructs were transformed into the *Agrobacterium tumefaciens* C58 strain. For histochemical GUS staining, constructs were transiently infiltrated into 4-week-old *rsi1* plants and after 48 hours post-infiltration, plants were challenged by MgCl_2_ (Mock) or *Pst*. 24 h post-pathogen infiltration, leaves were collected and stained with X-gluc solution and then destained using ethanol.

GUS activity was quantified by fluorometric enzymatic assay uning 4-methylumbelliferyl-β-D-glucuronide (MUG) as a substrate (Côté & Rutledge, 2003). Leaf samples were homogenized in liquid nitrogen in GUS extraction buffer containing 50 mM sodium phosphate buffer (pH 7.5), 0.1% (v/v) Triton X-100, 10 mM EDTA, 0.1% (w/v), 0.05% (v/v) β-Mercapto-ethanol and sodium lauroyl sarcosinate. After homogenization samples were centrifuged at 10000rpm for 15min. Supernatant was separated and total protein concentration was estimated using Bradford method. In 100 µL supernatant 100µL 4-MU was added following incubation in dark at 37°C for 10min. Reaction was stopped by incubating 100µL with 100µL 200mM Na_2_CO_3_. Fluorescence was measured in 96 well FLUOstar omega plate reader (BMG Labtech) at 360/460nm excitation and emission spectra. Remaining samples were kept in dark at 37°C and after 60 min, readings were taken again. For standards, known concentration of 4-MU (for MUG) and BSA (for Total protein estimation) was used.

### 2.3 Gene expression analysis

For RNA isolation, the guanidinium thiocyanate-phenol-chloroform method was used as described by Chomczynski and Sacchi (2006). For cDNA preparation, the iScript™ cDNA Synthesis Kit (#1708891, Bio-Rad, USA) was used, following the manufacturer’s protocol. Relative gene expression analysis was performed on a Bio-Rad CFX96 real-time PCR system (Bio-Rad, USA) using gene-specific primers. ACTIN2 (AT3G18780) was used as the housekeeping gene for normalization. For each experiment, three biological and two technical replicates were used (Giri et al., 2017; Singh et al., 2014).

### 2.4 Chromatin Immunoprecipitation Assay (ChIP)

ChIP was performed as described previously (Kumar et al., 2025; Saleh et al., 2008; Singh et al., 2013). 4-week old pRSI1:RSI-HA lines were treated with *Pst*-AvrRpt2 or Mock and after 3days, 4 gm systemic leaf samples were collected. For crosslinking of DNA-Protein crosslinking buffer (sucrose: 0.25 M; PIPES Ph 6.8: 15 mM; MgCl2: 5 mM; KCL: 60 mM; NaCl: 15 mM; CaCl2: 1mM; Triton X-100: 1%; protease inhibitor cocktail) was vacuum infiltration for 10 minutes. Crosslinking was stoped by adding 2.5 ml of 2 M Glycine (100mM final concentration) followed vacuum infiltration for 5 minutes at RT. samples were quickly frozen in liquid nitrogen and homogenized in cold 25 ml cold nuclei isolation buffer (sucrose: 0.25 M; PIPES Ph 6.8: 15 mM; MgCl2: 5 mM; KCL: 60 mM; NaCl: 15 mM; CaCl2: 1mM; Triton X-100: 1%; protease inhibitor cocktail). After filtering, homogenized samples were centrifuged at 13800 × g for 20 minutes at 4°c. The pellet was resuspended in nuclei lysis buffer [HEPES: 50mM (pH7.5); NaCl:150mM; EDTA: 1mM; SDS: 1%; Sodium deoxycholate: 0.1%; Triton X-100: 1%.], then sonicated and centrifuged to isolate chromatin with crosslinked proteins. After sonication samples were diluted 10 times and incubated with Protein A-Agarose beads for 2h at 4°C. Supernatant was separated which was incubated with the respective antibody (Anti-HA, Anti-H3K4me3, Anti-H3K4me2) at 4°C with gentle rotation and then Protein A-Agarose beads was added and incubated for 4-5 hours. Agarose beads were separated by centrifugation and washed according to the protocol. DNA-Protein Crosslinking was reversed by incubating samples with 5 M NaCl at 65°C overnight. After reverse cross-linking, proteins were degraded by Proteinase K and DNA was purified and resuspended in 50ul TE buffer which was used for qPCR.

## 3. Results and Discussion

### 3.1 Pathogen-induced *RSI1* expression is higher in *rsi1* mutant than WT plants

RSI1 codes for an epigenetic suppressor. However, whether RSI1 influences its own expression was not known. To investigate the role of RSI1 in its own expression, we monitored mRNA accumulation of RSI1 in systemic tissues of SAR-induced WT and *rsi1* plants. SAR was induced by infiltration with *Pseudomonas syringae* pv *tomato* DC3000 (*Pst*) carrying the *AvrRpt2* effector (*Pst-AvrRpt2*) at 10^7^ CFU/ml suspended in 10 mM MgCl_2_. The control (mock) group received only 10 mM MgCl_2_. *RSI1* transcript abundance was monitored by qPCR. As expected, WT plants showed enhanced expression of *RSI1* upon SAR induction compared to mock-treated plants (Fig. 1). However, to our surprise, we observed significantly higher levels of RSI1 expression in the *rsi1* mutant (Fig. 1). The results suggested the possibility of RSI1 functioning as a suppressor of its own expression.

**Fig. 1.**
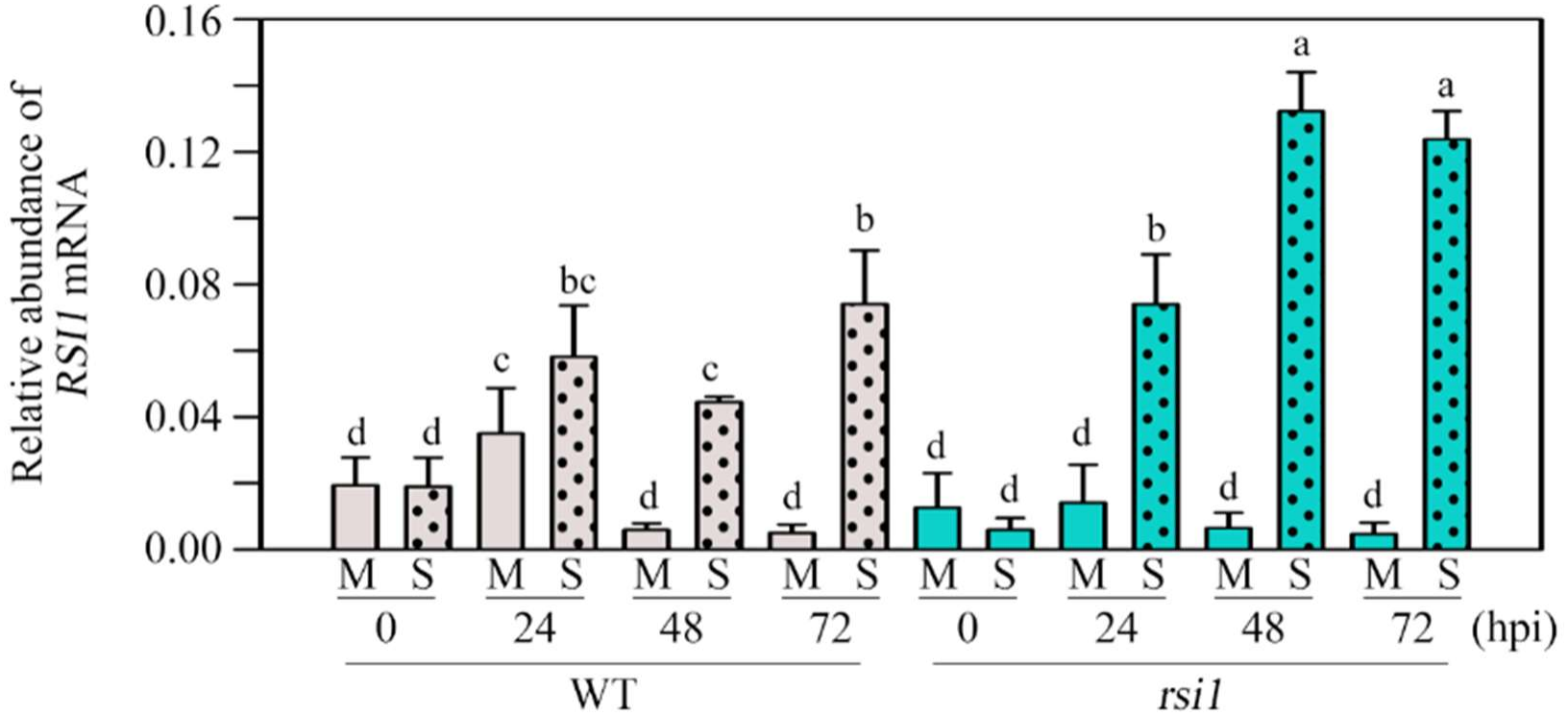
Abundance of *RSI1* mRNA in WT and *rsi1* plants. RSI1 mRNA abundance was analyzed in systemic tissue after Mock and SAR induction. Plants were infiltrated *Pst-AvrRpt2* (SAR) or Mock (10mM MgCl_2_), and systemic leaf tissues were collected at the mentioned hours after pathogen infiltration. RSI1 abundance was analyzed by qPCR. Each bar indicates the mean ± SD (n = 3). Different letters above the bars indicate values that are significantly different from each other as analyzed by one-way ANOVA (Holm-Sidak method, P < 0.001).

### 3.2 RSI1 suppress RSI1 promoter (pRSI1:GUS) activity

Increased level of RSI1 mRNA in *rsi1* mutant suggests that RSI1 suppresses its own expression. To confirm this finding, we have studied transient expression of pRSI1:GUS in the presence or absence of RSI1. Leaves of *rsi1* mutant plants were co-infiltrated with *Agrobacterium tumefaciens* cells containing the pRSI1:GUS construct and either p35S:RSI1-HA or vector control (p35S:HA). Two days after this infiltration same leaves were inoculated with *Pst-AvrRpt2* or 10 mM MgCl_2_ (mock). We harvested inoculated leaves after 24h of pathogen infiltration and performed histochemical GUS staining. In the mock-treated leaves, pRSI1:GUS expression was barely visible (Fig. 2A). We observed GUS expression by pRSI1:GUS construct at a high level after pathogen inoculation (Fig. 2A). Interestingly, when RSI1 was co-expressed with the p35S:RSI1-HA construct, GUS expression was severely reduced, suggesting a suppressive role of RSI1 on its own promoter activity. In contrast, the GUS staining levels from p35S:GUS were high in all conditions and were not affected by RSI1 expression. To validate the staining results, we quantified GUS activity by MUG assay. Results support the visible expression of GUS and clearly demonstrate that RSI1 expression significantly suppresses pRSI1:GUS activity (Fig. 2B).

**Fig. 2.**
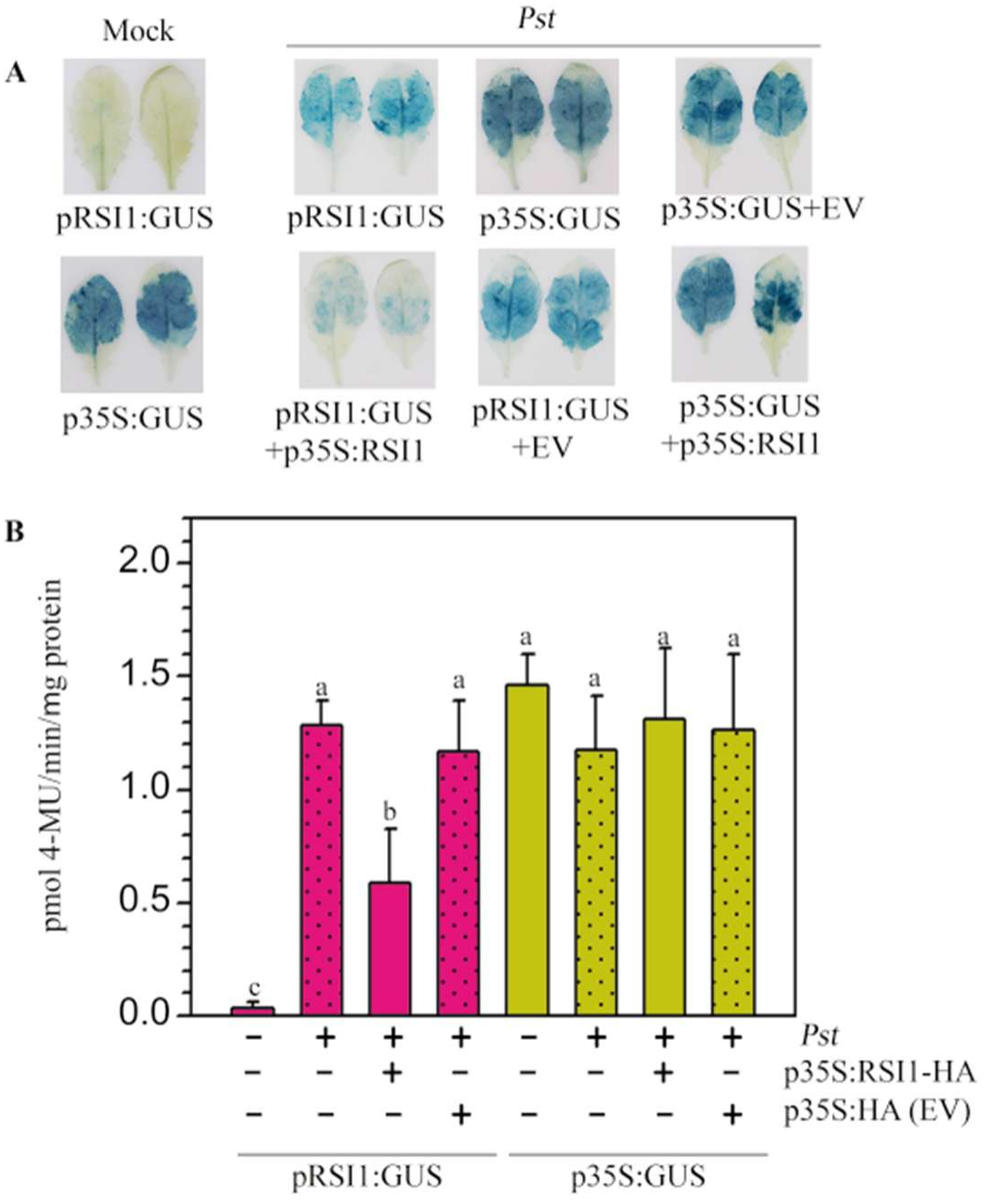
RSI1 suppresses pRSI1:GUS expression (A). GUS staining of *rsi1* leaves infiltrated with *Agrobacterium tumefaciens* containing pRSI1:GUS in the presence or absence of p35S:RSI1-HA. Empty vector (EV) was used as a control. Leaves were infiltrated by *Pst* or mock (10 mM MgCl_2_) **(B).** MUG assay to quantify GUS activity. Each bar represents the mean ± SD (n=3). Different letters above the bars indicate a significant difference in mean as determined by one-way ANOVA (Holm-Sidak method, P ≤ 0.001).

### 3.3 Pathogen-induced RSI1 expression requires GSTT2

RSI1 interacts with SANT domain-containing protein GSTT2 (Banday & Nandi, 2018). Function of GSTT2 is also required for the activation of SAR (Banday & Nandi, 2018). To study whether the functional SAR activation pathway is required for the pathogen-induced expression of RSI1, we monitored RSI1 mRNA accumulation in the *gstt2* mutant. We observed that RSI1 expression is induced by pathogen inoculation in local and systemic tissues only in WT plants but not in *gstt2* mutant (Fig. 3). The results indicate that intact SAR induction mechanism is required for the expression of RSI1 in local and systemic tissues.

**Fig. 3.**
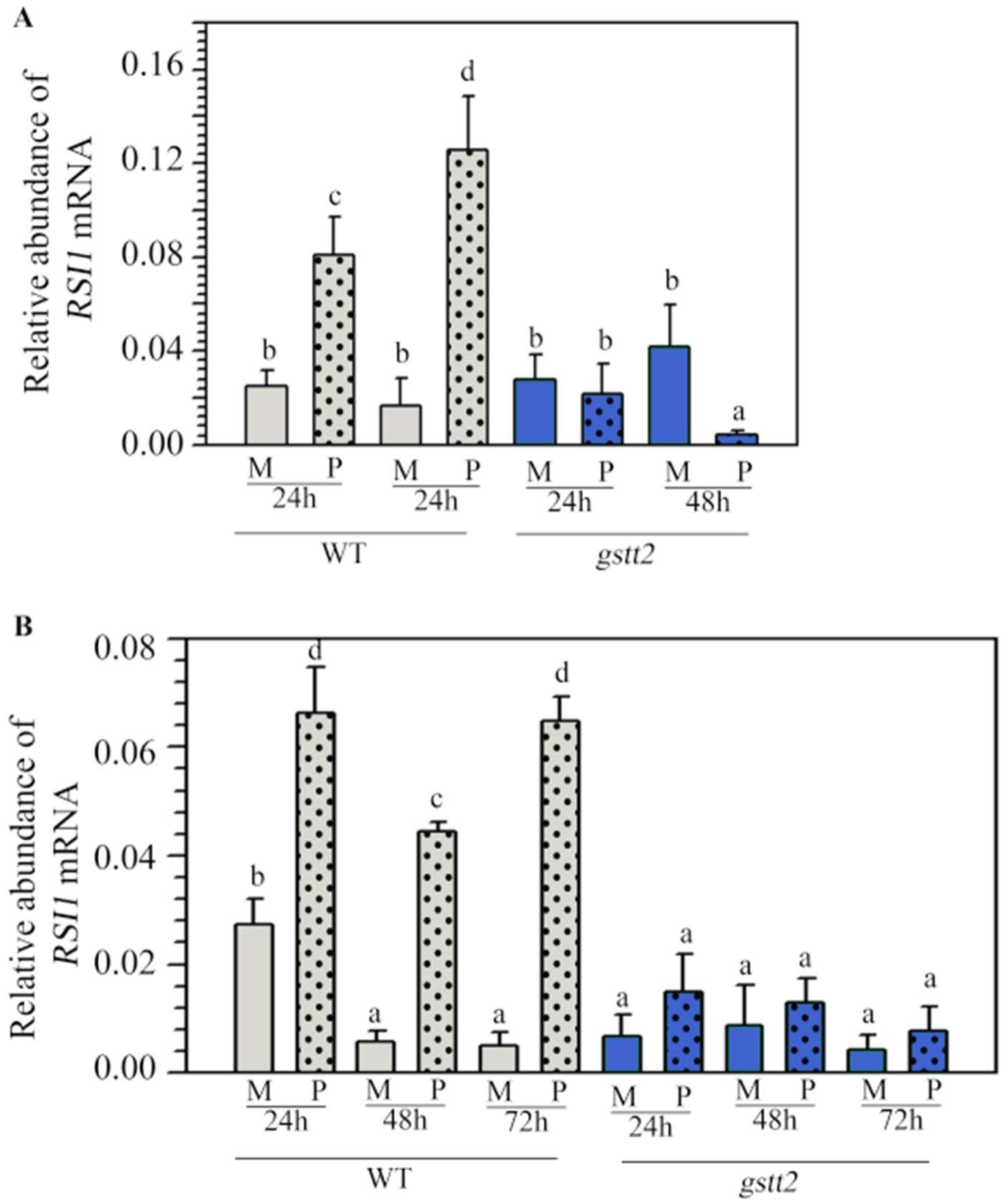
Relative abundance of RSI1 mRNA in WT and *gstt2* mutant. Abundance of *RSI1* transcript in WT and *gstt2* mutant was analyzed after Mock(M) and SAR induction(S) in local and systemic uninoculated tissue at indicated time points. RSI1 abundance was analyzes by qPCR. Each point indicates the mean SD (n=3). Different letters above the bars indicate values that are significantly different from each other as analyzed by one-way ANOVA (Holm-Sidak method, P < 0.001).

### 3.4 RSI1 physically associates with its own promoter

Experiments described above suggested the possible self-regulatory role of RSI1. To further confirm, we studied the physical association of *RSI1* at RSI1 locus. We used transgenic plants that express HA-tagged RSI1 under its native promoter (pRSI1:RSI1-HA) in *rsi1* mutant background (Kumar et al., 2025; Singh et al., 2023) and performed chromatin immunoprecipitation (ChIP) using anti-HA antibody. The abundance of four different regions in ChIP surrounding *RSI1* locus was studied by qPCR (Fig. 4A). We found the highest accumulation of RSI1-HA at the promoter-exon junction of the *RSI1* locus (Area C; Fig. 4B). The abundance of the amplicon in the precipitate reduced in both upstream (Area B) and downstream (Area D) of the transcription start site (TSS). No significant amplicons were detected in Area A. The results suggest that RSI1 specifically accumulated near the TSS in the RSI1 locus.

**Fig. 4.**
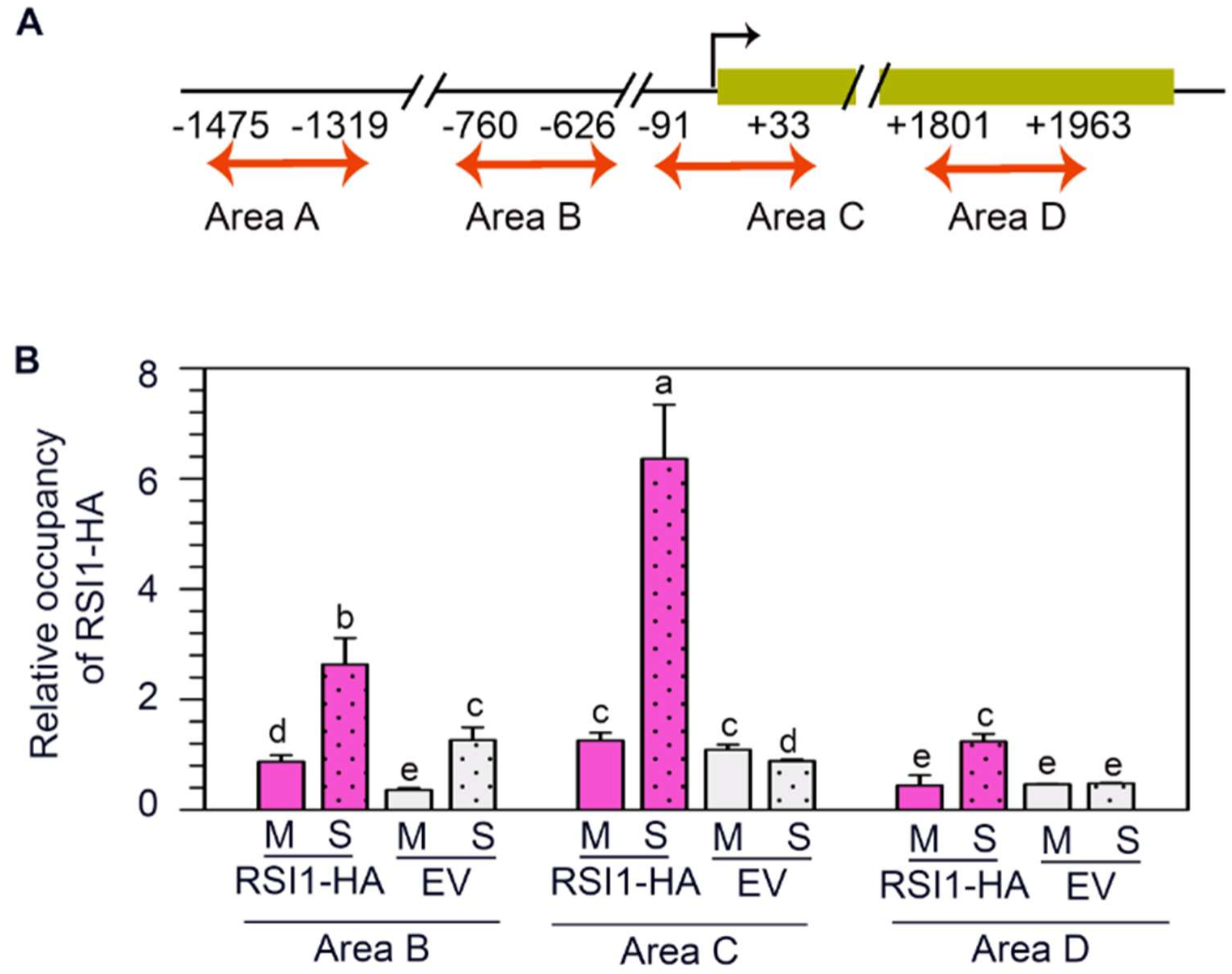
Occupancy of RSI1-HA at the RSI1 locus. **(A)** Diagram showing the areas used in the ChIP experiment at the RSI1 locus. **(B)** Relative abundance of RSI1-HA was precipitated by anti-HA antibody followed by qPCR at the RSI1 locus. Each bar represents the mean ± SD (n=3). Different letters above the bars indicate a significant difference in mean as determined by one-way ANOVA (Holm-Sidak method, P ≤ 0.001).

### 3.5 RSI1 demethylates H3K4me2 and H3K4me3 at *RSI1* locus

RSI1/FLD is a putative H3K4me2/me3 demethylase. To investigate whether RSI1 enzymatically removes methylation from H3K4me2/3 at the RSI1 locus, we performed ChIP experiment using anti-H3K4me2 and anti-H3K4me3 antibodies followed by qPCR at different regions of the RSI1 locus (Fig. 5A). We found that throughout the RSI1 locus, the abundance of H3K4me2 and H3K4me3 was higher in *rsi1* mutants than in WT plants (Fig 5B).

**Fig. 5.**
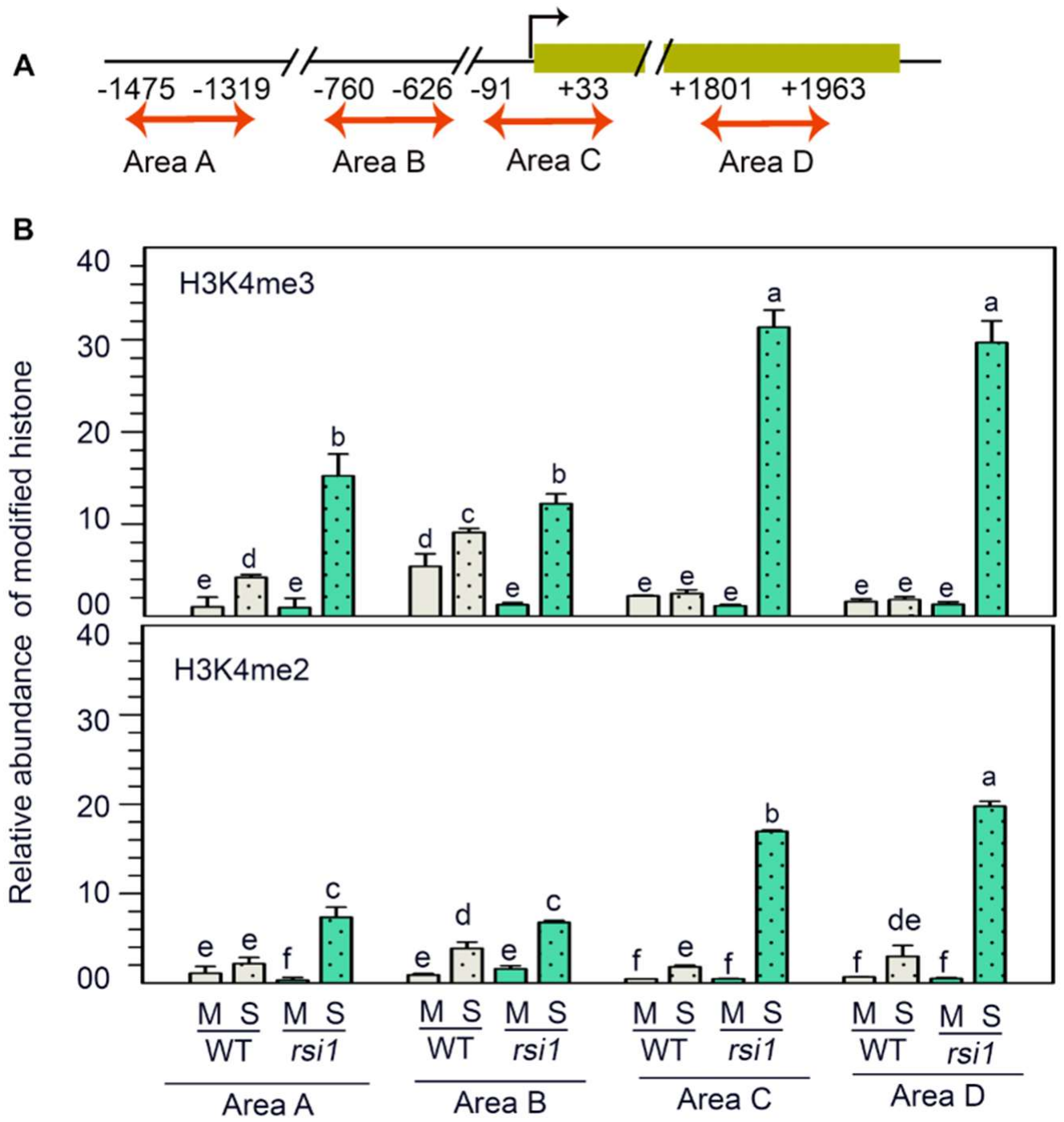
Levels of H3K4me3 and H3K4me2 at the RSI1 locus. **(A)**.Diagram showing areas used in ChIP experiment at the *RSI1* locus. **(B)**. Relative abundance of H3K4me3 and H3K4me2 on RSI1 locus. Each bar represents the mean ± SD (n=3). Different letters above the bars indicate significant differences in the mean as determined by one-way ANOVA (Holm-Sidak method, P ≤ 0.001).

## 4. Summary and Conclusion

The results described in this paper show that RSI1 expression of pathogen-inducible and intact SAR pathway is required for RSI1 expression both in local and systemic tissues upon pathogen inoculation. Intriguingly, the mRNA abundance of RSI1 is higher in the *rsi1* mutant than WT plants, which suggests that RSI1 negatively regulates its own expression. This hypothesis is experimentally confirmed by the physical association of RSI1 protein at the RSI1 gene near the transcription start site and by ChIP showing enhanced abundance of H3K4me2 and H3K4me3 throughout the RSI1 locus. Histone 3 with H3K4me2 and H3K4me3 modifications are associated with enhanced expression of the gene. Altogether, our data convincingly demonstrate that RSI1 negatively regulates its own expression. The possible reason for this negative regulation could be to keep SAR activation at an optimal level.

## Authorship contribution statement

**US, VP, RK, VS**-Experimental design, performing experiments, data analysis, writing manuscript draft. **AKN**-Conceptualization, experimental design, manuscript editing and reviewing. All authors approved the final manuscript.

## Data availability

Data is available from AKN upon reasonable request.

## Acknowledgement

This work was financially supported by the DBT grant (BT/PR52208/BSA/33/162/2024) to AKN. We also acknowledge DBT BUILDER grant (BT/INF/22/SP45382/2022) of SLS, and the PAIR grants (ANRF/PAIR/2025/000029/PAIR) JNU for additional support. US received JNU Non-NET Fellowship, VP received DST INSPIRE fellowship and RK received DBT fellowship during this work.

